# Structural basis and dynamics of Chikungunya alphavirus RNA capping by the nsP1 capping pores

**DOI:** 10.1101/2022.08.13.503841

**Authors:** Rhian Jones, Michael Homs, Nadia Rabat, Noelia Zamarreño, Rocio Arranz, Juan Reguera

## Abstract

Alphaviruses are emerging positive stranded RNA virus which replicate and transcribe their genomes in membranous organelles formed in the cell cytoplasm. The non-structural protein 1 (nsP1) is responsible for RNA capping and the gating of replication organelles by assembling into monotopic membrane-associated dodecameric pores (Jones R. et al. Nature 2021). The capping path is unique for Alphavirus; beginning with the N^7^ methylation of a GTP molecule, followed by the covalent linkage of a m^7^GMP group to a conserved histidine in nsP1 and the transfer of this cap structure to a diphosphate RNA (Ahola T. et al. PNAS 1995). Here we provide structural snapshots of different stages of the reaction pathway showing how nsP1 pores recognize the substrates of the methyl-transfer reaction, GTP and SAM, how it reaches a metastable post-methylation state with SAH and m^7^GTP in the active site, the subsequent covalent transfer of m^7^GMP to nsP1 and post-reaction conformational changes triggering the opening of the pore. In addition, we biochemically characterize the capping reaction, demonstrating specificity for the RNA substrate and the reversibility of the cap transfer resulting in decapping activity and the release of intermediates of the reaction. Our data identify the molecular determinants allowing each pathway transition, provide explanation for the need for the SAM methyl donor all along the pathway and new clues about the conformational rearrangements associated to the enzymatic activity of nsP1. Together our results set new ground for the structural and functional understanding of alphavirus RNA-capping and the design of antivirals.

**Significance statement:** Here we present biochemical and structural characterization of the capping pathway carried out by the Chikungunya virus nsP1 capping pores. We provide five Cryo-EM structures representative of the different steps of the reaction. These structures reveal the molecular determinants and dynamics associated with the alphavirus capping process. In addition, we biochemically show the RNA capping specificity and the reversibility of the reaction which allow nsP1 to cap and decap RNAs and to release intermediates of the reaction. These data provide a new biochemical clues on the enzymatic activity of nsP1 capping pores and a new structural landscape that will be instrumental for the design of effective antivirals targeting the viral RNA capping for blocking the infection.

## Introduction

Chikungunya virus (CHIKV) is an arbovirus transmitted to the human host by members of the *Aedes* mosquito family, causing infections that are characterized by fever, rashes, and debilitating joint pain. Although infections are rarely lethal and usually resolve in a few weeks, in some cases symptoms can persist for years (1), and the rapid spread of recent outbreaks has been a cause for concern (2). Alphaviruses such as CHIKV possess a positive sense single-stranded genome that must be replicated following release into the host cell. Genome replication occurs in membranous replication organelles (called “spherules”); invaginations that are derived from remodeling of the host cell membrane during infection (3). Each spherule houses a replication complex (RC) formed from four virally encoded non-structural proteins (nsPs) that function cooperatively in RNA synthesis (4). Within the RC, nsP4 is the RNA dependent RNA polymerase (5, 6), nsP2 has helicase activity (7) and proteolytically cleaves the nsPs from a viral polyprotein precursor (8, 9), and nsP3 has a role in recruitment of host factors to the spherule (10). NsP1 is the membrane anchor for the complex (11), forming dodecameric pores that associate monotopically with the membrane in the necks of the spherules to gate their entrance (12, 13). Enzymatically, nsP1 also has a role in addition of cap0 structures to the 5’ end of the positive sense viral RNAs (14–16). Cap structures, minimally formed from covalent linkage of an (m^7^GMP) moiety to the first nucleoside of the RNA via a 5’-5’ triphosphate bond (Cap0), are universally found in host mRNAs and are essential to the processing, stability, and translation of transcripts (17). In higher eukaryotes the 2’O ribose maybe further methylated (Cap1) along with internal bases of the mRNA. For viruses, capping of viral RNAs is thus often exploited as a means for hijacking the host translation machinery, and has an additional role in evasion of host innate immunity through preventing recognition of terminal RNA phosphates by cytosolic RIG-I and IFIT1 receptors (18). Alphaviral infection produces two viral RNA species capped by nsP1; the full length 11.8kbp positive sense genomic RNA (gRNA), and a sub-genomic RNA (sgRNA) of 4.3kbp that is transcribed from the second open reading frame (ORF2) at later stages of infection and encodes only the structural polyprotein (19, 20). Intriguingly, despite the central role of RNA capping in viral infection, recent studies suggest that not all alphaviral gRNAs packaged into virions are capped, and that uncapped RNAs may have an important role in modulating the host immune response to infection (21).

NsP1 directs cap synthesis via a mechanism that differs from the conserved capping pathways of most cellular and viral capping enzymes (15). In canonical pathways, a dedicated guanylyltransferase (GTase) enzyme transfers a GMP moiety from GTP to the 5’ phosphate of a diphosphate RNA (GpppA_N_). Methylation of the guanosine by a separate methyltransferase enzyme(s) yields the cap structure (m^7^GpppA_N_ for cap0). NsP1 possesses both N’7 methyltransferase and guanylyltransferase activity and reverses the order of these reactions. A methyl group must be transferred to GTP from a SAM substrate (forming m^7^GTP) prior to transfer of the m^7^GMP cap structure to the enzyme to form a covalent nsP1-cap0 intermediate on a conserved histidine (CHIKV-H37) (15). The cap is finally transferred to the 5’ terminal phosphate of a diphosphate RNA (m^7^GpppA_N_) (16). It is likely that the diphosphate RNA substrate is produced through the triphosphatase activity of nsP2, known to be specific to removal of γ-terminal phosphates (22).

Recent cryo EM structures of nsP1 capping pores (12, 13) and a partial RC (23) have provided important insights into nsP1 function, demonstrating that oligomerization into dodecamers is necessary for capping activity and driven by underlying interactions with the membrane. However, the structural basis for the non-canonical order of the pathway reactions is not understood, nor how the substrates are recognized or how many of the twelve sites within the ring can be active simultaneously. To address these questions, we provide a suite of cryoEM structures of nsP1 in complex with substrates from different steps of the capping pathway in addition to biochemical characterization of the RNA capping activity of the rings. The structures reveal that simultaneous binding of the SAM and GTP is necessary for optimal substrate positioning for guanylyltransferase activity, potentially providing a molecular rationale for why methylation precedes guanylation. We identify residues critical for substrate binding and residues playing a pivotal role in the different positioning of the guanosine triphosphate moiety during the capping process, including an arginine from a neighbor protomer in the pore (R275) which further explains the allosteric activation of nsP1 capping by pore formation. Our capping assays demonstrate that nsP1 exhibits sequence and structure specificity in capping of RNA substrates, suggesting that capping of the viral RNA occurs co-transcriptionally. We demonstrate that nsP1 is capable of releasing significant amounts of intermediates of the reaction and capable of decapping RNA substrates resulting in production of uncapped alphaviral RNAs. Finally, we show that the decapping reaction leads to a repositioning of regions important for the guanylyltransferase reaction in the nsP1 monomers, inducing opening of the pore. Together our results provide a large body of work for understanding the capping process of alphaviruses, revealing the molecular determinants and protein dynamics associated to this process. We discuss the implications of our findings on infection and host adaptation considering also recently reported structural findings on nsP1 pores (13).

## Results

### SAM binding to nsP1 is flexible in the absence of GTP

Each nsP1 protomer is built around a SAM dependent methyltransferase fold, with additional insertions (MBO loops 1 and 2) and extensions (RAMBO domain) that contribute to pore formation, oligomerization and membrane binding (Fig. S1A and B). In the context of the ring, the capping domains are located in the crown above the membrane, where a bilobal pocket in each protomer links a SAM binding site facing the exterior of the ring and a GTP binding site facing the interior (Fig. 1A and C). To better understand the initiating methylation step of GTP in the nsP1 the pathway, we solved the cryoEM structures of nsP1 in the presence of a 100 fold molar excess of SAM or GTP substrates (Table S1). Common to other N7-SAM dependent methyltransferases (N7 MTases), the SAM ligand binds in a pocket near the switch point of the Rossmann fold in the capping domain (Fig. 1B, Fig. S1A and B). The adenosine base and ribose sit within a cavity defined between loops β1-ηA above the SAM (including sequence motif ^63^DIG^65^ that is highly conserved in methyltransferases) and loop β2-αB below (including sequence motif ^89^DPER^92^) (see sequence alignment and legend in Fig. S2). The pocket is gated from the solvent exterior of the ring by loop αC-β4 above the zinc binding site. The SAM binding site is highly exposed to the solvent and largely defined by flexible loops (Fig. S1C and D).

**Figure 1.**
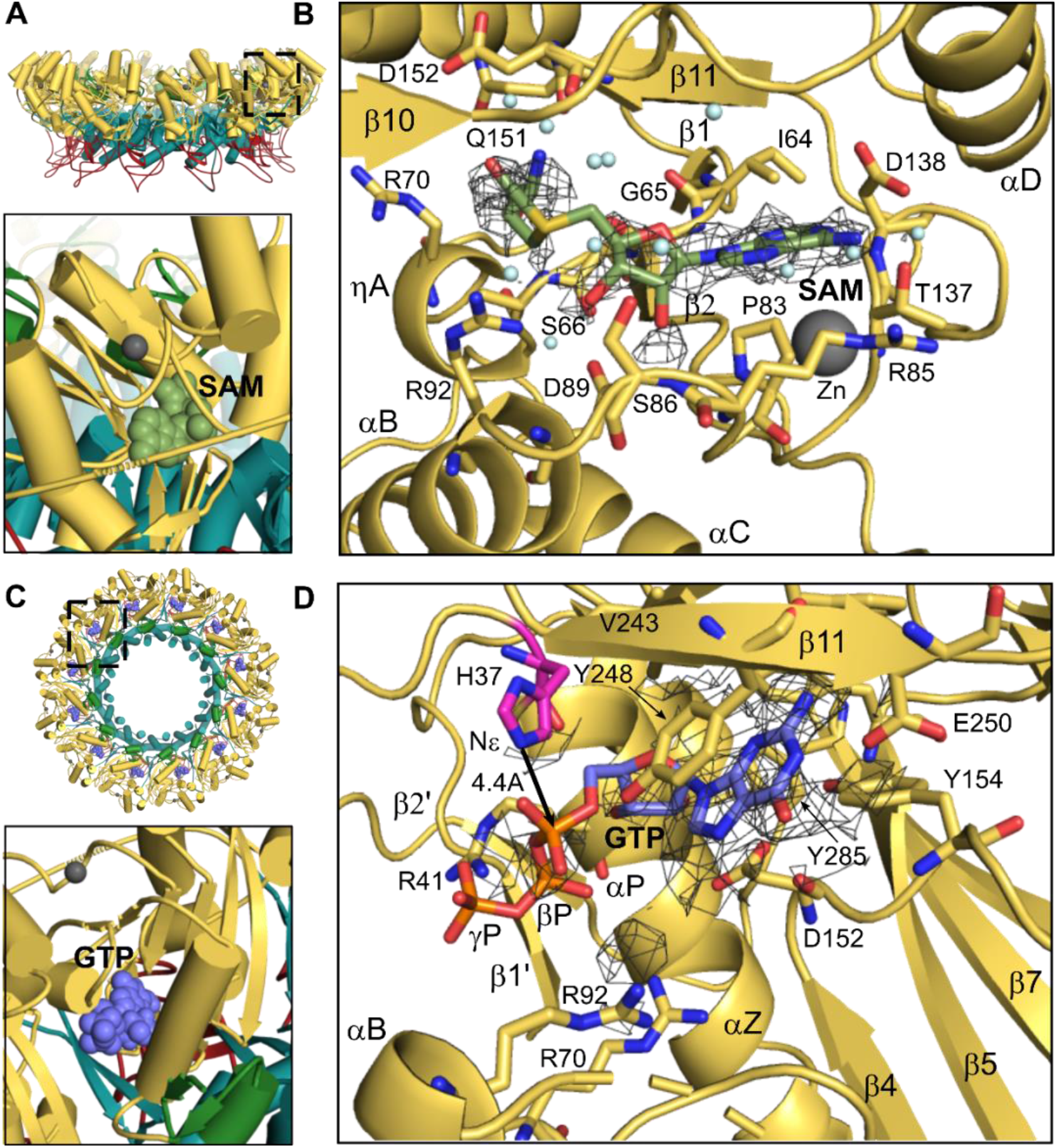
SAM and GTP substrate binding in nsP1 (methylation state). Panel A) Location of the SAM binding pocket in the SAM bound structure. The upper panel shows the nsP1 rings represented as a cartoon with the capping domain colored in yellow, the RAMBO domains in teal, and the membrane binding loops in red. The SAM binding site (boxed) faces the exterior of the ring. The lower panel shows the SAM molecule as green spheres in one monomer of the nsP1 dodecamer. Panel B) Molecular details of the SAM substrate binding site. The amino acid residues interacting with SAM are represented by sticks and labeled, in addition to the secondary structures that define the site. A detail of the cryoEM map for the SAM ligand is shown as a gray mesh contoured at sigma level 2. The map for the ligand becomes ambiguous beyond the purine and ribose base and many residues around the ligand appear to be flexible. Panel C) Top view representation of the ring as in A. Location of one of the GTP binding sites is indicated by a square in the nsP1 dodecamer. The bottom panel shows the position of the GTP ligand (purple balls) facing the ring interior. Panel D) Details of the GTP binding pocket shown as in B with the cryoEM map for the ligand contoured at sigma level 2. GTP-interacting residues from nsP1 are labelled, and the distance from Nε of the catalytic His37 residue (highlighted in magenta) to the alpha phosphate is indicated. The density for the phosphate moieties is only observed at higher contour levels. All Figures were made in Pymol.

The density for the SAM ligand is not fully defined in the binding pocket, where in maps reconstructed with symmetry, only density for the base and ribose moieties is clearly visible (Fig. 1B). In comparison with the apo form of nsP1, neighbouring helices αB and C are very poorly defined and overall local resolution for this region is worse compared to the apo and other bound states (Fig. S3). Contacts made to the SAM ligand are primarily weak van der Waal’s (VdW) interactions, involving residues I64-A67 from the DIG motif and P83, R85, T137 and D138 to the base; S86, D89 and R92 from the DPER motif to the ribose and residues R70, G151 and D152 to the methionine (Fig. 1B and Table S2). The SAM base forms a hydrogen bond (H-bond) between N6 and the side chain of residue T137 in nsP1, which maintains a second H-bond with R85. Both residues are conservatively mutated among alphaviruses (Fig. S2). Interestingly, D138, which is highly conserved in other N7 MTases and typically confers SAM binding specificity through an H-bond to the N6 amine, is barely defined in the density of the disordered αC-β4 loop and too distant to contact the amine (∼4Å).

The purine base is bound in the *anti*-conformation to the ribose, where the 2’ and 3’ hydroxyl groups of the ribose form H-bond to the side chains of D89 of helix αB. Mutagenesis studies have identified both D89 and R92 as important for SAM binding (24). Although the density for the methionine is quite poorly resolved, its position replaces the side chain of R70 in the apo form of nsP1, which is also poorly defined in the structure. The methionine is surrounded by residues G65, S66 and A67, and Q151 and D152, which are within H bonding distance to the amine group of the methionine.

To determine whether differences in protomer conformation or substrate occupancy could underlie the poor definition of the SAM binding site in the maps, we performed focused classification following symmetry expansion with a mask centered on the capping domain (Fig. S4). We could identify four classes by focused classification. In 20.6 % of the particles the SAM site was empty (class 3), despite the high molar excess of SAM added to the protein. For a second class (class 1) including another 18% of particles, only density corresponding to the purine and ribose of the SAM is visible but the remainder of the active site is clearly defined. In a third class (Class 2), including 22.9% of the particles the ribose and methionine are also clearly defined, but density for helices αC and αB above the SAM binding site were barely visible.

Taken all together, these data suggest that in the absence of GTP, SAM is not stably bound, and does not satisfy some of the contacts required for correct positioning of the substrate for methyl transfer. The correlation between the presence of SAM in the active site and the disordering of helixes αC and αB suggest that binding of the SAM alone also induces significant destabilization of part of the nsP1 capping domain. This could explain why apo nsP1 does not co-purify with significant amounts of SAM or SAH, as is observed for various other SAM dependent N7 MTases (DENV NS5 (25, 26), human RNMT(27)), and contrasts with N7 MTases that require SAH/SAM cofactors for stabilization and crystallization (vaccinia MTase) (28).

### GTP binding occurs in a deeper pocket relative to other N7-MTases and requires occupation of the adjacent SAM pocket

The GTP pocket in the capping domain of nsP1 is defined by the tip of strand β4, loop β4-αD and helix αZ, where loops αC-β4 and β4-αD communicate with the adjacent SAM site (Fig. 1C and D). Strand β11 of the beta sheet insertion in the capping domain forms a lid over the pocket. The guanosine base fits in an hydrophobic pocket defined by residues D152, Y248, F241 (β10-11 lid), Y154 (β4-αD loop), and F178 (β5). Base stacking occurs in between D152 and Y248, where the latter residue changes rotamer relative to the SAM bound and apo structures to become planar with the guanosine. E250 of strand β11 forms H-bonds to the N1 and N2 of the guanosine base, and the carboxylate group of residue D152 (β4-αD loop) maintains a network of H-bonds with GTP-O6 and N7 through an intermediate water molecule. Both contacts are conserved in other N7 MTases and confer specificity for methylation of a GTP substrate over adenine, where the N1 position is unprotonated and interaction with the E250 carboxylate would be unfavourable (29).

However, relative to other N7 MTases, the GTP pocket is deeper in nsP1, and the guanosine base and ribose are bound further in the fold (Fig. S1D). The overall effect of this deeper binding pose compared to canonical N7 MTases is to align the alpha phosphate of the GTP with H37, a catalytic residue identified to form a phosphoramide bond in the m^7^GMP-nsP1 covalent intermediate (15, 16). However, the position of the GTP alpha phosphate is still at 4.8Å from the histidine side chain Nε, thus too distant to undergo nucleophilic attack. Density for the secondary structures defining the GTP site is well defined compared to the SAM site, implying greater rigidity of the binding site (Fig. S3). However, as for the SAM ligand, only the base and ribose of the GTP are clearly defined in the maps, suggesting that the phosphates are flexibly bound. The side chains of the positively charged arginine residues lining the path for the phosphates (R92, R70 and R41) are also disordered, with the exception of R41, which becomes ordered relative to the apo and SAM bound structures and forms H-bonds to the poorly defined β and γ phosphates. Residues C82-89 that connect to the adjacent SAM site are also disordered, and increasing contour levels in the map reveals continuous density beyond N7 of the GTP that aligns with the binding path for SAM/SAH ligands but could not be assigned, as if the SAM binding pocket was base-promiscuous to a certain extent and GTP was non-specifically occupying the site. In conclusion, the structure clearly shows that many of the residues essential for GTase transfer (see next section) are flexible with only GTP bound, and that further stabilization of the GTP is required for subsequent methyl transfer and GTase reactions. The superposition of both SAM and GTP bound structures shows that the GTP N7 is apically positioned for an in-line SN2 nucleophilic attack and at 2.8 Å of the SAM methyl group (Fig. S5A). The arrangement of SAM and GTP bound alone to the active site are thus representative of the pre-methylation state of the reaction.

### Simultaneous binding of SAH and m^7^GTP induces a metastable conformation of the active site

To investigate guanylation of nsP1, the second step of the reaction, we acquired structures of the protein with SAH and m^7^GTP in the absence of magnesium, a cofactor necessary for nsP1 GTase activity. We found in our structure, as expected, that both ligands occupy the active site in a state corresponding to the post-methylation reaction but prior to the m^7^GMP transfer to the nsP1. The structure of nsP1 in the post methylation state shows no overall conformational changes with respect to the SAM and GTP bound structures. However, the definition of the ligand and SAM binding site densities drastically improves, suggesting that the ligands are more stably bound together and that this induces an ordering of the active site (Fig. 2A).

**Figure 2.**
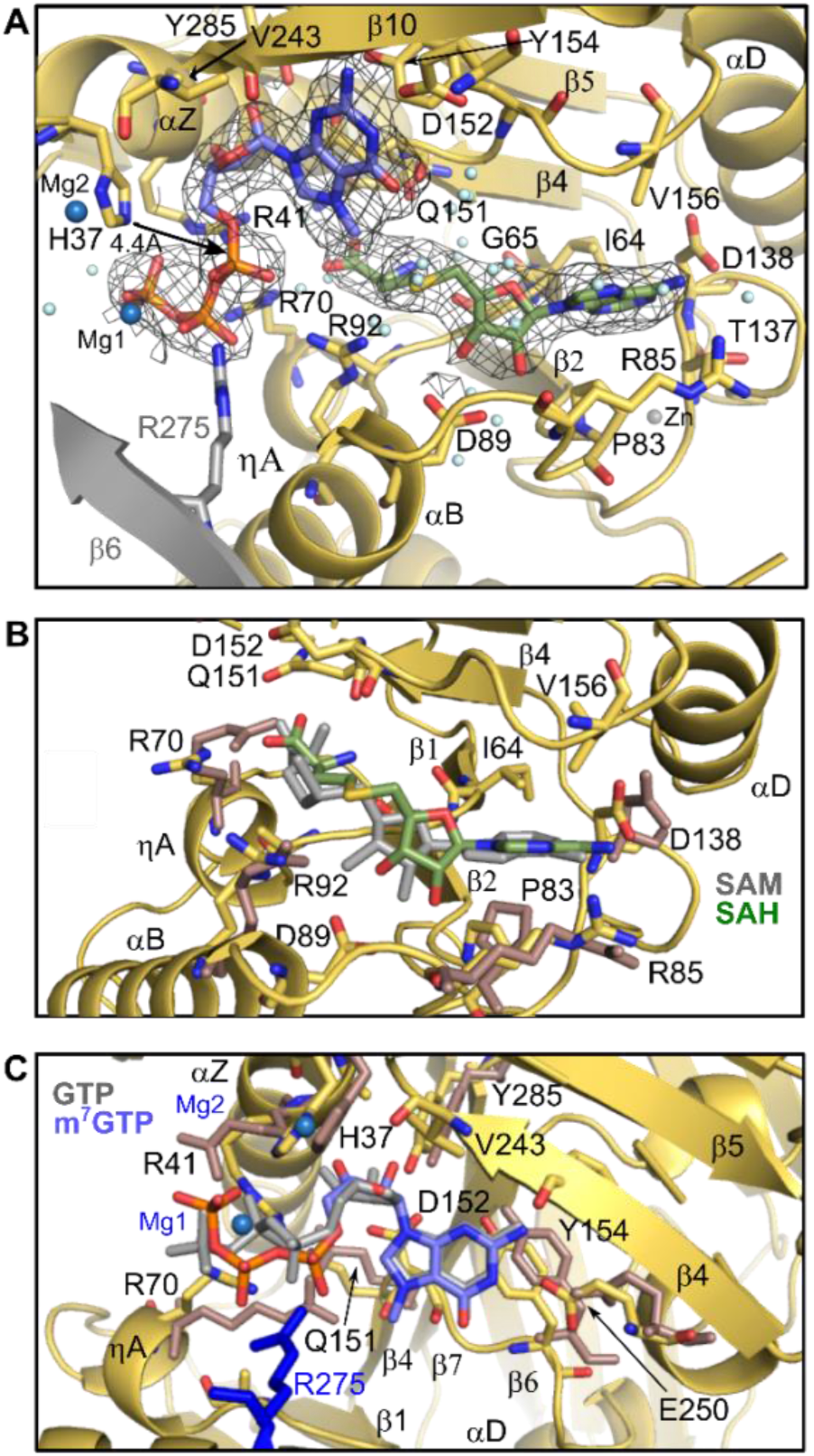
SAH and m^7^GTP ligand binding in nsP1 (guanylation state). Panel A) Detail of the nsP1 active site from the SAH and m^7^GTP bound structure. Density for the ligands from the cryoEM maps is contoured at 2 sigma and is substantially improved relative to the structures with the ligands bound separately, particularly for the cysteine moiety of the SAH and the phosphates of the m^7^GTP. However, the phosphate residues are still sub-optimally positioned for nucleophilic attack. Residues and secondary structures from the neighboring protomer that form contacts to the m^7^GTP are colored grey. Panel B) Overlay of the SAH bound site (SAH substrate in green) with the SAM bound structure (ligand in grey). Residues that have changed conformer relative to the SAM bound structure are colored pink. Panel C) Overlay of the m^7^GTP binding site (m^7^GTP in lilac) with the GTP bound structure (GTP ligand in grey). Side chains that have changed rotamer conformation are colored pink as in C. Figures were generated in Pymol.

Many contacts formed to the SAH purine and ribose are shared with the SAM structure, (G65, P83, D89, D138 and V156) (Fig. 2B and Table S2). However, there is a slight rotation in the SAH ribose and base (Fig. 2B), resulting in an ordering of the αC-β4 loop and bringing the side chain of D138 within H-bonding distance of N6 of the SAH base. Additional contacts are made to the methionine, which is now clearly anchored within the cavity. The methionine carboxyl forms VdW / weak H-bonds with the backbone of G65 and side chain of R70. The side chain of R92 becomes ordered and forms a contact to the methionine sulphur, held in position by an interaction with the N^7^ methyl group of the m^7^GTP (Fig. S5B). The methyl and sulphur moieties are still apically positioned as for an in-line SN2 nucleophilic attack at 3.4 Å of distance, slightly longer than in the pre-methylation state (2.8 Å). This suggests that there is minimal relocation of the substrates following the first methylation reaction and R92 appears to be important for stabilizing the sulphur leaving group in the methylation reaction.

There is significant movement of the m^7^GTP phosphates and an ordering of the surrounding arginine residues (Fig. 2C) relative to the GTP bound structure. R41 forms a new hydrogen bond to the oxygen bridging the alpha and beta phosphate, and R70 moves to bridge the beta and gamma phosphates. R92 and R275 of the neighbouring protomer move in to directly coordinate the alpha phosphate aligned with H37 (Fig. 2C and S5B). This contact provides direct evidence that oligomerization is required to complete the GTP substrate binding site, explaining why nsP1 monomers are inactive for GTase activity (12).

However, from analysis of the structure, it is immediately clear that the substrate is not correctly positioned for nucleophilic attack by the catalytic histidine for guanylation. The histidine Nε is still 4.4Å away from the alpha phosphate (Fig. 2A) and the beta and gamma phosphate leaving group are not apically positioned to the nucleophile for formation of a pentavalent transition state.

Surprisingly, attempts to capture the m^7^GMP covalently bound intermediate by adding 2mM MgCl_2_ and incubating for 2 hours yielded exactly the same configuration of the nsP1 active site substrates with two new densities attributable to Mg^2+^ metal ions. Mg1 appears to coordinate Pβ and Pγ O2B and O3G respectively and a second Mg2 atom appears to coordinate the catalytic H37 and D36 sidechain at 2 Å and 2.3 Å distance of each respectively (Fig 2A and S5B). In this conformation, the Pα is still too far from H37 for the nucleophilic attack and there is no density for a magnesium ion in proximity to the alpha phosphate that is usually necessary to increase its’ electrophilicity to promote attack. The structure suggests that Mg2, coordinated with the catalytic histidine and D36, could be playing this catalytic role by coordinating with Pα after repositioning of the m7GTP phosphates to be ready for the nucleophilic attack. In conclusion, the presence of m^7^GTP and SAH in the active site results in a metastable conformation that cannot transition directly to the covalent transfer of m^7^GMP to the catalytic H37, possibly constituting a regulated checkpoint of the capping reaction during infection.

### Structural basis of the guanyltransfer reaction

We finally obtained electron microscopy maps corresponding to the nsP1-m^7^GMP covalent complex by incubation of the nsP1 protein with the m^7^GTP and SAH substrates and RNA, suggesting that the presence of RNA is able to bypass the metastable state and stimulate the guanylation reaction (Fig. 3A). Here the position of the guanosine base moves as the ribose and alpha phosphate rotate by 56°, bringing the Pα close enough to H37 for establishing a covalent bond (Fig. 3B). As a consequence of the phosphate repositioning, residue R41 is now closer to the alpha phosphate and a new H-bond is formed between the ribose and S44 sidechain (Fig. S5C). In parallel Y248 is now too far from the alpha phosphate for hydrogen bonding, and sidechains of residues R92 and R275 of the neighboring protomer are no longer visible in the maps (Fig 3B). Intriguingly, the SAH substrate remains bound in the SAM binding pocket now extensively contacting by VdW interactions the m^7^GMP moiety as a consequence of its rotation (from 3 to 14 contacts, see Table S2). There is no density for the pyrophosphate product, suggesting it leaves with the magnesium ions. The H-bonds of the base with E250, and the ribose with Y285 are maintained despite the rotation of the base. E250, Y285 and R41 appear to be the pivotal residues on which the GMP moiety turns for approaching the catalytic histidine, R41 being next to the Pα transfer reaction site. The conformation of R41 is supported by the stacking of R70, which reaches the active configuration only after SAM/SAH binding. This indicates that both residues, highly conserved in alphavirus nsP1 (Fig. S2), are central for the switch between the methylation and guanylylation steps of the nsP1 capping reaction in coordination with R92 and R275. In all the steps, the binding of SAM or SAH is necessary for the proper configuration of the active site for the binding of the different GTP derived intermediates, whether by direct VdW contacts between methionine and base or, more importantly, by residues sharing contacts with both GTP and SAH such as D152, R70 or R92.

**Figure 3.**
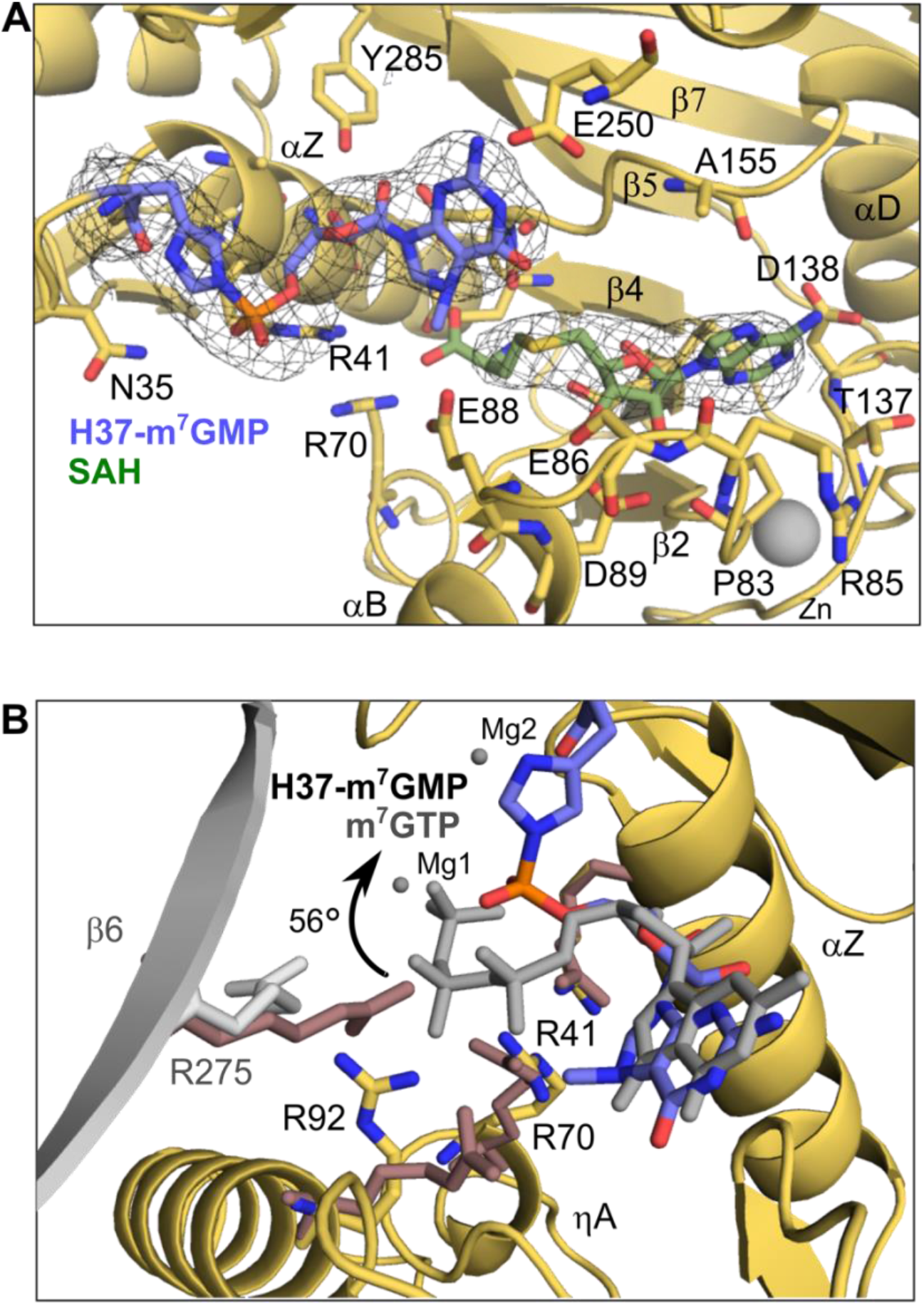
The covalent nsP1-GMP cap structure. Panel A) Detail of the active site, with density contoured at 2 sigma showing the covalent bond formed between catalytic histidine 37 and the m^7^GMP cap. Panel B) Overlay of the covalent m^7^GMP-H37 moiety and the m^7^GTP from m^7^GTP/SAH bound structure (m^7^GTP ligand in grey). The image shows a rotation of the alpha phosphate by 56° towards the catalytic histidine. Arginine residues around the catalytic site from the m^7^GTP structure are colored pink and overlaid with those from the m^7^GMP covalent structure, where the β6 strand from the neighbouring protomer is coloured in grey. R70, R95 and R275 from the neighboring protomer all change side chain rotamer, whilst R41 remains fixed as the residue that the nucleotide pivots around. Figures were generated in Pymol.

### The NsP1 RNA capping reaction is structure and sequence specific and reversible

To investigate the final step of the reaction, cap transfer to different RNA substrates was followed using ^32^αP labelled GTP and autoradiography. Alphaviral gRNAs and sgRNAs contain highly conserved and stable stem loop structures in the 5’UTR just downstream of the cap (Fig. 4A and Fig. S6), that may be important for transcription from the minus strand sequence during the replication cycle. To investigate the role of such stem loop structures in capping, we compared capping of a sequence corresponding to the first 27 nucleotides of the CHIKV gRNA sequence preserving the first stem-loop structure, and capping of a 15-nucleotide substrate that truncates this loop (Fig. 4A and B). Only the 15mer was appreciably capped when di-or tri-phosphorylated at the 5’ end, but not with a free 5’ hydroxyl group (Fig. 4A). The similarity in capping activity for di and tri-phosphorylated RNA suggest that the same capped forms are achieved, and implies the possibility of a concomitant triphosphate hydrolysis during the guanyl transfer reaction. The 27-nucleotide long CHIKV RNA was not capped in any of the phosphorylated states (Fig 4A, Fig S7), and following prolonged incubation times, only lower molecular weight RNAs of similar length to the 15mer were capped, presumably the products of partial RNA degradation or residual transcription products (Fig. S7). This indicates that the RNA binding cavity is too narrow to accommodate a double stranded RNA stem loop structure.

**Figure 4.**
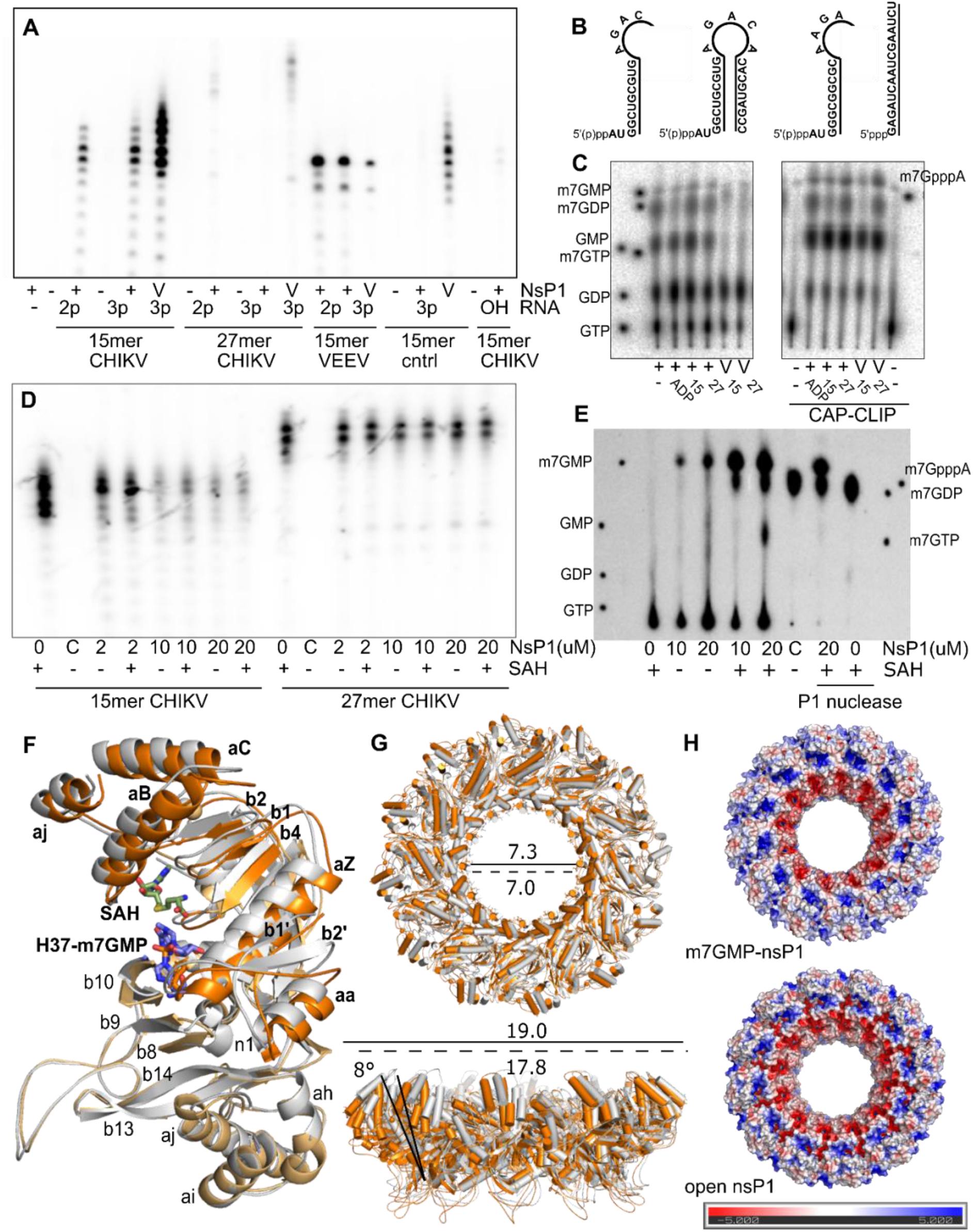
NsP1 RNA capping is reversible and sequence and structure specific. Panel A) Autoradiograph of a 20% urea gel with the products of the capping reactions performed in the presence of ^32^PαGTP and SAM. 2 μM NsP1 was incubated with 5μM RNA (15mer CHIKV, 27mer CHIKV, 15mer VEEV and a control RNA beginning GAG) with a diphosphate (2p) or triphosphate (3p) at the 5’ end, or with a free hydroxyl (OH). The vaccinia capping enzyme (V) was used as a positive control. Panel B) Schematic of the different RNA substrates synthesised via *in vitro transcription* for this study, corresponding to the 15 and 27 nucleotide 5’UTR of CHIKV (strain S27), the 15 nucleotide 5’UTR of VEEV, and a 16 nucleotide control RNA sequence corresponding to the 5’UTR of Crimean Congo Hemorrhagic fever virus (CCHFV). RNA substrates were synthesized with a diphosphate or triphosphate 5’ end, and a 15mer for the CHIKV sequence with a free hydroxyl group at the 5’ end was purchased commercially. 27mer and 15mers with a cap0 structure were generated using a commercially available vaccinia capping enzyme for decapping experiments. Panel C) Autoradiograph of TLC for selected capping reactions following digestion with P1 nuclease, where ADP was also used as a control substrate. A CAP-Clip enzyme was used to confirm for the presence of a bona fide cap structure. Panel D) Autoradiograph of 20% urea PAGE gel of purified capped RNAs (estimated final concentration at 2 μM) incubated in the presence of increasing concentrations of nsP1 in the absence or presence of 2mM SAH. A decapping enzyme from *S. pombe* was used as a positive control reaction (labelled C). Panel E) Autoradiograph of TLC plate showing the products of selected decapping reactions. A reaction with and without nsP1 was digested with P1 nuclease as a control. Panel E) Structural overlay of an nsP1 monomer from the closed form of the rings (m^7^GMP covalently capped structure in grey) and the open form obtained from the decapping reaction (orange). There is a substantial movement in the first 130 residues of the capping domain and C-terminal helix, coloured in deep orange and labelled in bold typeface. Panel F) The structures of the open and closed ring forms overlaid, colored as in E. A tilting of the capping domains 8° away from the central pore results in an overall expansion of the pores in the rings obtained from the decapping reaction. Dimensions of the ring and central aperture are indicated with a solid line for the open conformation and a dashed line for the closed conformation. Panel G) Difference in charge distribution in the rings for the closed and open forms.

When investigating sequence specificity, nsP1 was also able to cap a 15-nucleotide long RNA with a sequence derived from the 5’UTR of Venezuelan Equine Encephalitis virus (VEEV), a related new world alphavirus where the first four nucleotides of the sequence are conserved with CHIKV. However, nsP1 was unable to cap an unrelated control RNA of the same size with a different initiating 5’ RNA sequence (beginning GAG) (Fig 4A). Despite considerable sequence variability in the 5’UTR sequences of alphaviruses, 3 of the first 4 nucleotides (the first AU and fourth G) in gRNAs and sgRNAs are highly conserved between Old and New World viruses and possibly the determinants of specificity (Fig. S7). Thus, the capping reaction is specific for Alphavirus RNAs short enough to prevent the formation of secondary structures. This results are consistent with the recent identification of the U as the nucleotide recognized by the nsP1 through specific interactions with residues at the N-terminal extension of the capping domain (30).

Digestion with P1 nuclease and analysis of the products by thin layer chromatography (TLC) was used to confirm for the presence of a *bona fide* cap m^7^GpppA structure (Fig. 4C). Intriguingly, in presence or absence of RNA, where GTP and SAM were present in the reaction, species migrating as a cap structures (m^7^GpppA) were consistently observed in TLC. This suggests that nsP1 has the capacity to cap the GTP nucleotide or any contaminating GDP, something that has already been observed for nsP1 from VEEV (31). In addition, m^7^GTP, the product of the methyl transfer reaction, as well as m^7^GDP and m^7^GMP, which are not products on the reaction pathway, were identified. The nucleotide capping activity of nsP1 could potentially alter the GTP/m^7^GTP homeostasis of the cell and potentially affect many cellular GTP dependent metabolic processes.

The m^7^GMP is a product that could only be released from hydrolysis of the m^7^GTP-nsP1 covalent complex or from loss of the cap from the RNA in a decapping reaction. To test for the latter possibility, we labelled the triphosphate 27mer and 15mer CHIKV RNAs with a ^32^αP cap0 structure using the vaccinia capping enzyme. RNAs were isopropanol precipitated to remove excess nucleotides and incubated with increasing ratios of nsP1 in the presence and absence of SAH (Fig. 4D). Whilst signal for the 27mer was unchanged, a decrease in radioactivity was observed for the 15mer as the concentration of nsP1 was increased, whether in the absence or presence of SAH. TLC analysis of the reaction products with and without P1 nuclease digestion confirmed that this corresponded mainly to loss of m^7^GMP, and not m^7^GDP as in the commercial *S. pombe* decapping enzyme control reaction (Fig. 4E). This confirm that the nsP1 capping reaction is reversible, conferring the enzyme with decapping activity.

Together these results suggest that capping of the alphaviral RNA most likely occurs co-transcriptionally prior to folding of the conserved SL1 loop. This would protect the viral RNA from decapping as the loop folds during synthesis, whilst other Alphavirus-like cellular mRNAs could be potentially decapped.

### The decappping reaction induces conformational changes in nsP1 and opens the capping pores

Capping and decapping are two directions of the same reaction. In order to investigate the structure of the pores after the capping/decapping reaction we analysed by cryo-EM the structure of nsP1 pores incubated with a cap0 11-nucleotides-long CHIKV RNA. All the previously described structures in this article have the same overall conformation of the pore. However, after the decapping reaction we could distinguish two different 3D classes, one similar to the previous structures and a second with significant changes in the first 130 residues and the C-terminal alpha helix k of nsP1 (Fig. 4F). Electron density for the m^7^GMP base was found in the GTP binding site in both classes. The nsP1 protomers appear tilted outwards 8° with respect to the equatorial axis of the ring (Fig. 4G). The pore opening results on an increase of the inner aperture of 3 Å (from 70 to 73 Å) and the outer diameter of 12 Å (from 178 Å to 190 Å) and a change in the surface charges distribution concomitant with a projection of the active site towards the top of the ring (Fig. 4G and 4H). This conformation resembles the structure of nsP1 pores when expressed in mammalian cells (see discussion)(13). Thus, we can conclude that the decapping reaction induces a motion in the ring resulting in an opening of the pore aperture.

## Discussion

Although the nsP1 capping mechanism has been well characterized enzymatically over the years, the structural basis for the non-canonical order of the pathway has remained elusive. The highly symmetrical capping pores of chikungunya present an opportunity for detailed structural characterization of the nsP1 capping pathway via analysis of cryo-EM structures that represent the different stages of the pathway.

The structures show that although many of the contacts formed to the SAM/SAH and GTP/m^7^GTP substrates are conserved with other N7 MTases, there are significant differences in the configuration of the active sites that may have allowed the protein to evolve additional GTase activity. Notably, the GTP binds in a deeper pocket in nsP1, aligning the alpha phosphate with the catalytic histidine for cap transfer. It appears that there is minimal movement in the positions of the substrates between the methylation and guanylation reactions, but several key side chains change positions to mediate the transition between methytransferase and guanyltransferase activity. Simultaneous binding of the substrates appears to be necessary for engaging these residues and for correct positioning of the substrates within the active site. NsP1 is unable to robustly form a covalent complex with GTP or m^7^GTP in the absence of SAM/SAH (12, 15), and we demonstrate that when bound alone, the GTP and SAM substrates exhibit substantial flexibility beyond the purine base. In comparison, the SAH and m^7^GTP structures bound together are more stably anchored within the cavity by more stable contacts made to the nsP1 protein. The displacement of R70 by the methionine moiety of the SAM ligand appears to be essential for the positioning of the surrounding residues and the GTP phosphates in the active site, and in transfer of the cap to H37 the guanosine base conformation is positioned by stacking with the SAH molecule. Overall, these structural details explain why methylation must precede guanylation in the capping pathway of nsP1.

Our study also provides insights into the sequence and structural preferences of nsP1 for RNA substrates. We show that RNA capping is reversible and suggest that capping occurs co-transcriptionally prior to folding of the SL1 loop, which may have a protective role in preventing decapping of the RNA following its’ formation in addition to preventing recognition of the cap0 structure by host IFIT1 (18). Recent studies have suggested that only a small percentage of alphaviral gRNAs packaged into virions are capped (21). Although our understanding of the roles of these RNAs in infection is still in its infancy, it has been demonstrated that increasing the capping activity of nsP1 in the context of a virus is detrimental to SINV infection, suggesting that capping activity must be finely tuned (32). Finally, future research is required to address if nsP1 has the capacity to decap cellular mRNA substrates, which are mainly modified with a cap1 structure in higher eukaryotes, contributing to host translational shut-down. Certain viruses, including *poxviridae*, encode viral decapping enzymes that are expressed at later stages of infection and remove cap structures on host mRNAs to prevent their recognition by eIF4e at the ribosome (33). Such strategies require that the viral RNA be translated via an alternative ribosomal recognition mechanism, such as an IRES in *poxviridae*. It has been reported that a conserved stem loop structure downstream of the initial AUG codon of the alphaviral sgRNA may promote translation independently of eIF4G in an infection context (34, 35). The sequence specificity that we find in our capping experiments suggests that nsP1 could target specific cellular mRNAs, initiating AUG. To our knowledge, there are no precedents of enzymes with both capping and decapping activities. Both activities would need to be highly regulated in the context of the RC.

While this manuscript was in preparation Zhang et al(30) have reported an independent study presenting structures of some of states of the capping reaction provided here with nsP1 capping pores produced in mammalian cells. The expression and purification of capping pores in mammalian cells delivers complexes with an expanded pore conformation. When we express nsP1 in insect cells and purify the pores with the same detergents and conditions used in their study (see methods) we consistently observe a contracted conformation of nsP1 pores. Thus, the expression system and not the purification protocol determines the nsP1 conformation, whether due to different lipid compositions of the inner plasma membrane or different components in the cytoplasm.

The methyl transfer and first guanylyltransfer reactions occur without significant conformational changes in the nsP1 structure in both mammalian and insect cells derived pores. Here, we show how the post-decapping state induces motions that trigger an opening in the pore, similar to the open form of nsP1 complexes expressed in mammalian cells. This open form increases surface exposure of the RNA binding pockets, with a concomitant redistribution of surface changes along a path leading to the internal pore. All of these changes can potentially induce changes in the full replication complex that may determine different stages of RC functioning in the late and early steps of infection. Since differences in capping pore conformations exist depending on the expression system, these could result in different behaviours of the RC in the host reflecting host-adaptation of the replication machinery.

Our structure of the nsP1 complex with SAH and m^7^GTP shows nsP1 in a post methylation state and metastable initial state of the guanylyltransferase reaction. This unreacted state is found in the expanded and contracted forms of the nsP1 pores (purified from mammalian and insect cells respectively), and is thus not dependent on ring conformation. Covalent cap formation is only observed following incubation with RNA, whether in the form of a capped 12mer or with a non-reactive 27-nucleotide-long RNA in the presence of m^7^GTP and SAH. The RNA is thus able to trigger the reaction, even if not stably binding to the complex by, for instance, changing the exposed arginine distribution that hold the m^7^GTP in the postmethylation state. Indeed, the hydrogen bonds maintained between R92 and R275 of the neighbouring monomer and the gamma phosphate of the m^7^GTP must be broken to allow for the rotation of the alpha phosphate observed in the nsP1 m^7^GMP covalently linked structure, bringing the phosphate group within attacking distance of H37 and apically positioning the pyrophosphate leaving group. Intriguingly, despite the difference in overall fold, such a metastable state has also been described for other GTases (36), where an opening and closing of the active site induces phosphate relocation and GMP transfer from a GTP substrate.

Interestingly when we incubate nsP1 in presence of SAH and m^7^GTP we can see by western blot or radioactivity, in denaturing conditions, the covalent link by an antibody specific for m^7^GMP. This is indeed a well-established test for guanylyltransferase activity (12),(16). These data suggest that rather than a particular conformation of the pores (expanded or contracted) it appears that some event is required to trigger the reaction, whether the presence of RNA (physiological conditions) or by treating the sample with denaturant agents. Our data strongly suggest that in the context of the RC the covalent transfer of m^7^GTP to the H37 could be a regulated checkpoint of the capping reaction.

In conclusion, the different structural snapshots of the capping reaction presented here describe in detail the individual role of the residues involved in substrate recognition and N7 methylation and guanyltransferase reactions. The results show that SAM and GTP substrate binding are interdependent and essential for ordering of the active site to allow all steps of the reaction. We identify residues R70 and R41, not present in other conventional MTases, as main players for the GTP binding and transfer of the Cap0 to His 37. These results provide a mechanistic explanation for the peculiar alphavirus capping pathway characterized biochemically over decades, paving the way for future research on understanding alphavirus RNA capping and the structure-based design of antivirals against alphavirus infections.

## Methods

### NsP1 protein purification

NsP1 was expressed in Hi5 cells (Thermo Fisher) as outlined in Jones et al. (12) using baculovirus technology. Protein samples of nsP1 with GTP or SAM substrates were purified as described in Jones et al(12), in fos-choline12 detergent. For the SAH and m^7^GTP and m^7^GMP complexes, this method was adapted to obtain single rings using the protocol of Zhang et al. (13), where the solubilization step was performed with 1% DDM and samples were exchanged into 0.01% GDN. Briefly, following recovery of the membranes from lysed cells by ultracentrifugation at 100,000g, membranes were resuspended at 100mg/ml in 35mM tris, 200mM NaCl, 1mM TCEP and 5% glycerol with 1% DDM for 2 hours at 4°C. The soluble fraction recovered post centrifugation at 100,000g was applied to Ni-NTA resin in batch (1ml of resin per gram of solubilised membrane) and washed with 10 column volumes of wash buffer containing GDN to exchange the detergent (35mM tris pH 7.6, 200mM NaCl, 1mM TCEP, 5% glycerol, 40mM imidazole, 0.01% GDN). Samples were eluted in elution buffer (35mM tris pH 7.6, 200mM NaCl, 1mM TCEP, 5% glycerol, 300mM imidazole, 0.01% GDN) and concentrated with a centrifugal concentrator with a 100kDa MW cut off prior to application to a Superose6 10/30 column in gel filtration buffer (25mM HEPES pH 7.6, 150mM NaCl, 1mM TCEP, 0.01% GDN). For both fos-choline and GDN purified samples, the central peak fraction was selected for cryoEM.

### Sample preparation for cryoEM

For the SAM and GTP nsP1 samples, 0.3mg/ml of nsP1 was incubated with 0.5mM of each substrate in gel filtration buffer (25mM tris pH7.6, 150mM NaCl, 1mM TCEP, 0.065% fos-choline 12). 3μl of each sample was applied to Quantifoil R2.2 Copper Rhodium grids (mesh size 300) with a homemade carbon coating after glow-discharging for 1 minute at 100mA. Samples were vitrified in a Vitrobot mark IV using blot force 0 for 3 seconds at 25°C and 95% humidity.

For the SAH and m^7^GTP, nsP1 was incubated at 0.2mg/ml in gel filtration buffer (25mM HEPES pH 7.6, 150mM NaCl, 1mM TCEP, 0.01% GDN) with 0.5mM SAH and 0.5mM m^7^GTP in gel filtration buffer supplanted with 2mM MgCl_2_, and incubated for 2 hours at 30°C prior to direct application to an EM grid and freezing. For the formation of the m^7^GMP intermediate, the same protocol was followed but a 10 fold excess of the CHIKV 27mer RNA was added to the reaction just before freezing. 3μl of sample was applied to Quantifoil R2.2 gold grids (mesh size 300) that had been coated with a homemade film of graphene oxide after glow discharging for 10s at 100mA. Samples were vitrified in a vitrobot mark IV using blot force -3 for 3 seconds at 25°C and 95% humidity.

To verify that the detergent was not altering the conformation or flexibility of the active site in the protein, for the SAH and m^7^GTP complex a dataset was also collected for a sample purified in fos-choline12 for direct comparison, where the maps showed no significant differences.

### Cryo-EM data collection

With the exception of the m^7^GMP nsP1 covalent complex, all final datasets were collected on a Krios at CM01 of the ESRF at 300kV equipped with a post column LS/97 energy filter (Gatan), slit width 20eV. For the SAM and GTP datasets, images were acquired with a K2 summit camera in counting mode at a nominal magnification of 165,000, (corresponding to a sampling rate of 0.827Å or 1.06Å per pixel, see table S1) across a defocus range of 1 to 2.5um. For the SAM dataset 4500 movies were recorded with a dose rate of 7.2 e-per pixel per s for an exposure time of 4s distributed over 40 frames, yielding a total accumulated does of 42.4e-per Å^2^. The GTP dataset was recorded with a dose rate of 15.6 e-per pixel per s for an exposure time of 3.4s distributed over 40 frames, yielding a total accumulated does of 42e-per Å^2^. 3077 movies for the SAH and m^7^GTP dataset were recorded with a K3 camera operating in super-resolution mode, with a super-resolution pixel size of 0.42Å and nominal magnification of 105,000. Total dose was 38e-distributed over 40 frames, with a dose rate of 14.9e-per pixel per s for an exposure time of 1.85s.

The m^7^GMP dataset was recorded on a TALOS Artica microscope operating at 200kV (Instruct platform, CNB Madrid). 773 movies were recorded with a Falcon III camera operating in counting mode, at a nominal magnification of 120,000 and corresponding pixel size of 0.855Å per pixel. Accumulated dose was 32e-per Å^2^ in 38s, distributed over 60 frames with a dose rate of 0.73e-/pix/s.

### Cryo-EM data processing

Datasets were analysed in parallel in Relion (version 3.0) (37) and cryoSPARC (38). Frame alignment and correction for beam induced motion was performed in MotionCorr2 (39) using patch alignment, and CTF correction was performed with CTFFind4(40) from non-dose weighted micrographs. Images with poor ice quality, excessive astigmatism or with no Thon rings beyond 5Å in Fourier power spectra were discarded from further processing. Particle picking was performed from dose-weighted micrographs using Warp (41) or Relion’s template matching method, using templates that had been generated from an initial round of picking and 2D classification. Particles were extracted with a box size of 300-360 pixels, and were binned twice for initial processing. 2D classification in Relion or cryoSPARC was used for removal of bad particles, and an *ab initio* model was generated from these particles without imposing symmetry. 3D classification performed with the *ab initio* models was used for separation of single and double rings for datasets purified in fos-choline detergent, but otherwise did not reveal any differences in conformations or occupancy state between rings. Classification was performed with a soft spherical mask of 290, and with coarse alignment sampling (7.5°). For each dataset, the best 3D class was used for auto-refinement in Relion or non-uniform refinement in cryoSPARC in c1, following re-extraction of particles to the original pixel size. As no significant differences in monomers were observed, refinements were repeated imposing c12 symmetry and with masking. Maps were sharpened using post-processing in relion. Masks used for refinement and sharpening were generated through filtering of the reconstructed volume to 15Å, and through extended the mask by 3 pixels and adding a soft edge of 3 pixels.

To look for differences between protomers in rings, focused classifications of the capping domain was performed following symmetry expansion of the particles. Particle sets from c12 refinement were expanded using relion_symmetry_expand to align all protomers. A mask placed around a single capping domain was used to perform signal subtraction on the remainder of the images, where subtracted particles were reboxed to 90 pixels on a region centered around the mask coordinates. The capping domain mask was generated as outlined above, using an .mrc map generated from a single capping domain using the mol2mapcommand in chimera from the nsP1 PDB. The subtracted images were reconstructed without alignment to generate a reference for 3D classification. 3D classification was performed without alignment using between 3-8 classes for robustness, with a T value of 25. Resulting maps were sharpened suing Phenix autosharpen map (42). All models were built into the cryo-EM maps using the nsP1 PDB structure 6Z0V (12), and were subjected to iterative rounds of refinement and model building in Phenix(42) and Coot (43).

### Synthesis and purification of RNA substrates

RNA substrates corresponding to the first 27 or 15 nucleotides of the CHIKV genome (strain S27) were synthesized by *in vitro* transcription, with a Type II promoter to yield an AUG starting codon. To obtain substrates with a 5’ diphosphate, a five-fold excess of ADP was added to the other nucleotides for synthesis. RNAs were resolved on an 8M urea 20% acrylamide gel and extracted using sodium acetate and isopropanol precipitation. Post washing of the pellets with 70% ethanol, RNAs were resuspended in water and stored at - 20°C until use. Sample quality was assessed by 8M-urea PAGE, analysis of A^260^/A^280^ and A^230^/A^260^ ratios, and an aliquot was digested with P1 nuclease (NEB #M0660S) and analysed by LC-MS to identify the phosphorylation state of the nucleotide at the 5’ end.

### RNA capping assays

For RNA capping assays, 2μM nsP1 was incubated with 100 μM SAM, 1 μM α^32^PGTP (to have a final specific activity of 0.1uCurie/ul in a reaction) and 5μM RNA at 30°C for 2 hours in capping buffer (50mM HEPES pH 7.6, 50mM KCl, 5mM DTT and 2mM MgCl_2_). Transfer of the m^7^GMP cap to the RNA via 8M urea PAGE (20% gel) using autoradiography. The commercially available vaccinia virus capping system (NEB #M2080S) was used as a positive control.

Thin layer chromatography (TLC) was used to confirm for the presence of the cap structure and identify other lower molecular weight products. 5ul of each reaction was digested with P1 nuclease and treated with proteinase K (NEB #P8107S), prior to application to a TLC membrane (Machery-Nagel) pre-activated in absolute ethanol. Samples were pre-migrated in water and then transferred to 0.65M Li_2_SO_4_ or 1M (NH_4_)_2_SO_4_ as a mobile phase. The membrane was dried and visualized using autoradiography, comparing to migration standards.

### RNA decapping assays

To produce capped RNAs, 20μM of CHIKV 27mer or 15mer RNA was incubated with vaccinia capping enzyme (NEB #M2080S) in the presence of 1mM SAM and α^32^PGTP (to have 0.3uCi/ul specific activity in the final reaction) for 1 hour at 37°C. The enzyme was heat inactivated at 75°C for 2 minutes and then removed with 1μl of Strataclean resin (Agilent). Capped RNAs were precipitated using 2M ammonium acetate and isopropanol to remove any residual nucleotides, and the pellet washed twice with 70% ethanol prior to resuspension in the same volume of H_2_O used for the initial reaction. The resuspended RNA was incubated at a final estimated concentration of 2μM with increasing molar ratios of nsP1 (from 1:1-10:1) for 2 hours at 30°C in capping buffer with or without 100μM SAH. RNA concentration was calculated assuming 50% recovery from the precipitation reactions. RNA incubated in the absence of nsP1 and with *S. pombe* mRNA decapping enzyme (NEB #M0608S) were used as negative and positive controls. Loss of the cap from the RNA was followed with autoradioagraphy and 20% acrylamide urea PAGE or TLC, as described above. For decapping assays, TLC was performed with or without P1 nuclease digestion.

## Acknowledgments

We thank Eaazhisai Kandiah and the ESRF for the access to the Titan Krios at the ESRF CM01 through the French BAG, and Daouda Traore as a local contact. We thank Jaime Martin-Benito and the EM platform at the CNB-CSIC for granting us access to cryo-EM equipment through a technical support contract, Javier Chinchon and Roberto Melero at the CNB, CSIC and to Denis Ptchelkine and the electron microscopy platform at the AFMB for technical assistance. This work has been supported by the Bettencourt Shueller Fondation and the ATIP-avenir program (CNRS/INSERM). R.J. was supported by Instruct ERIC with a short-term fellowship (APPID 1717) to access the Cryo-EM facility at the CNB, CSIC (VID 31525 and 31528). This work made use of ScipionCloud and the resources provided by the IFCA cloud site, which have been partially supported by the project ‘European Open Science Cloud - Expanding Capacities by building Capabilities’ (EOSC-SYNERGY), funded by the RI Horizon 2020 Program of the European Commission under Grant Agreement No. 857647. Experiments performed at the PBSIM facility at the AFMB were supported by the French Infrastructure for Integrated Structural Biology grant ANR-10-INSB-05-01.

The structures have been deposited in the PDB with accession codes 8AOX (SAM bound), 8AOV (GTP bound), 8AOW (m^7^GTP/SAH bound), and 8APX (m^7^GMP covalent complex). The corresponding maps have been deposited in the EMDB with accession codes EMD-15555, EMD-15553, EMD-15554, and EMD-15578.

