## Supplementary information for "Structural basis and dynamics of Chikungunya alphavirus RNA capping by the nsP1 capping pores"

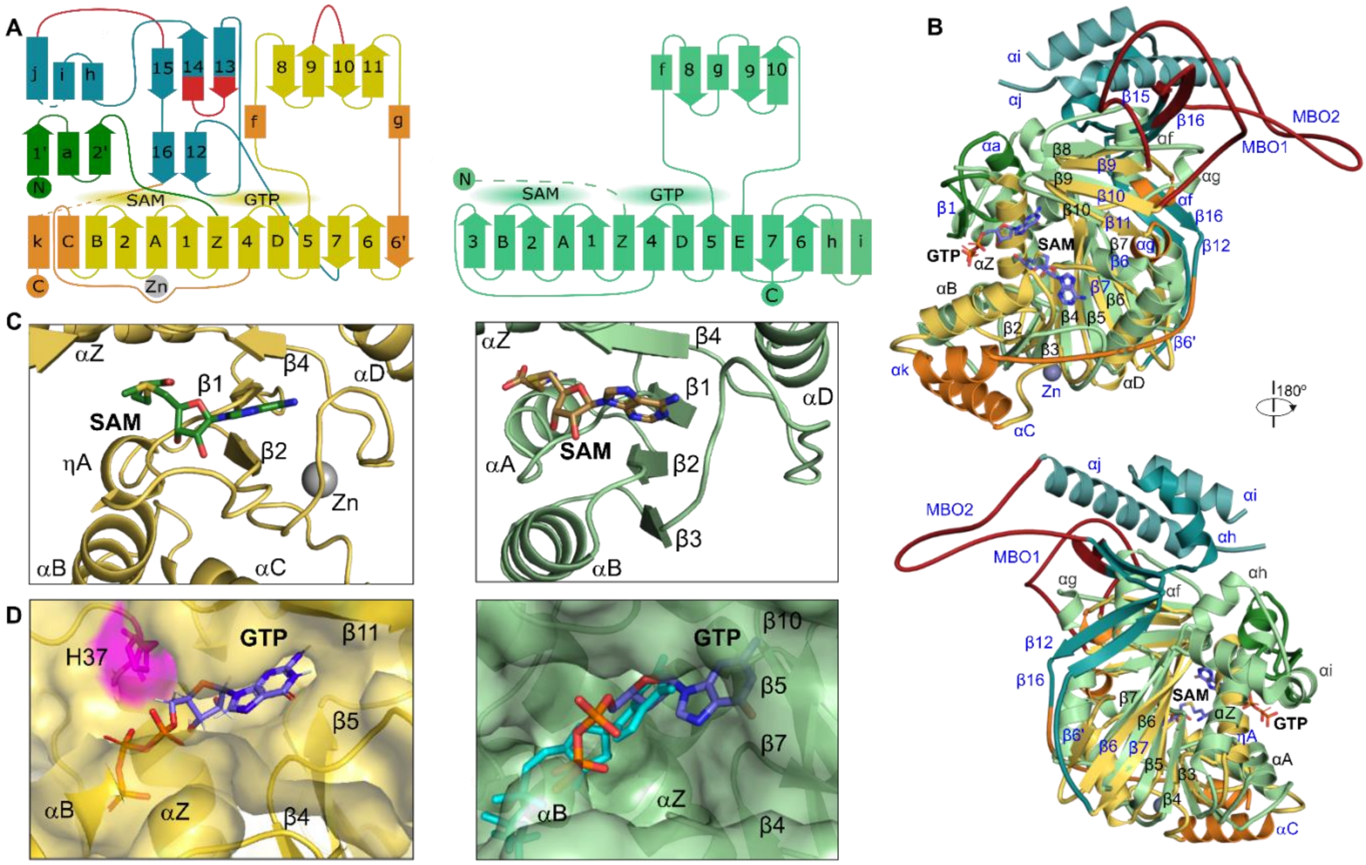

**Figure S1: Structural comparison of an nsP1 protomer and canonical methyltransferase (Ecm1)**(1)**.** A) Topology comparison of the two proteins. Left, the conserved capping domain of nsP1 is in yellow, with insertions in the capping domain relative to Ecm1 colored orange. The nsP1 N-terminal domain is colored dark green, and the C-terminal ring aperture membrane binding oligomerisation (RAMBO) domain is in teal with membrane binding loops (MBO1 and 2) in red. The Ecm1 protein is represented in light green on the right. B) Overlay of the structures, colored as in A. Secondary structural elements that are unique to nsP1 or that overlay with Ecm1 but differ in their numbering are labelled in blue. C) Comparison of the nsP1 (left) and Ecm1 (right) SAM binding pockets, colored as in A. The loss of strand β3, shortening of strand β1 and extended loop αC-β4 means that the site in nsP1 is mostly defined by flexible loop regions. D) Comparison of the nsP1 (left) and Ecm1 (right) GTP binding pockets, with the proteins colored as in A. The binding pocket is deeper in nsP1 to align the alpha phosphate with catalytic H37. In Ecm1, the GTP (cyan) binds further out, and a structural overlay with the nsP1 bound GTP (purple) demonstrates that residues from strands β5 and β7 would exclude this binding pose.

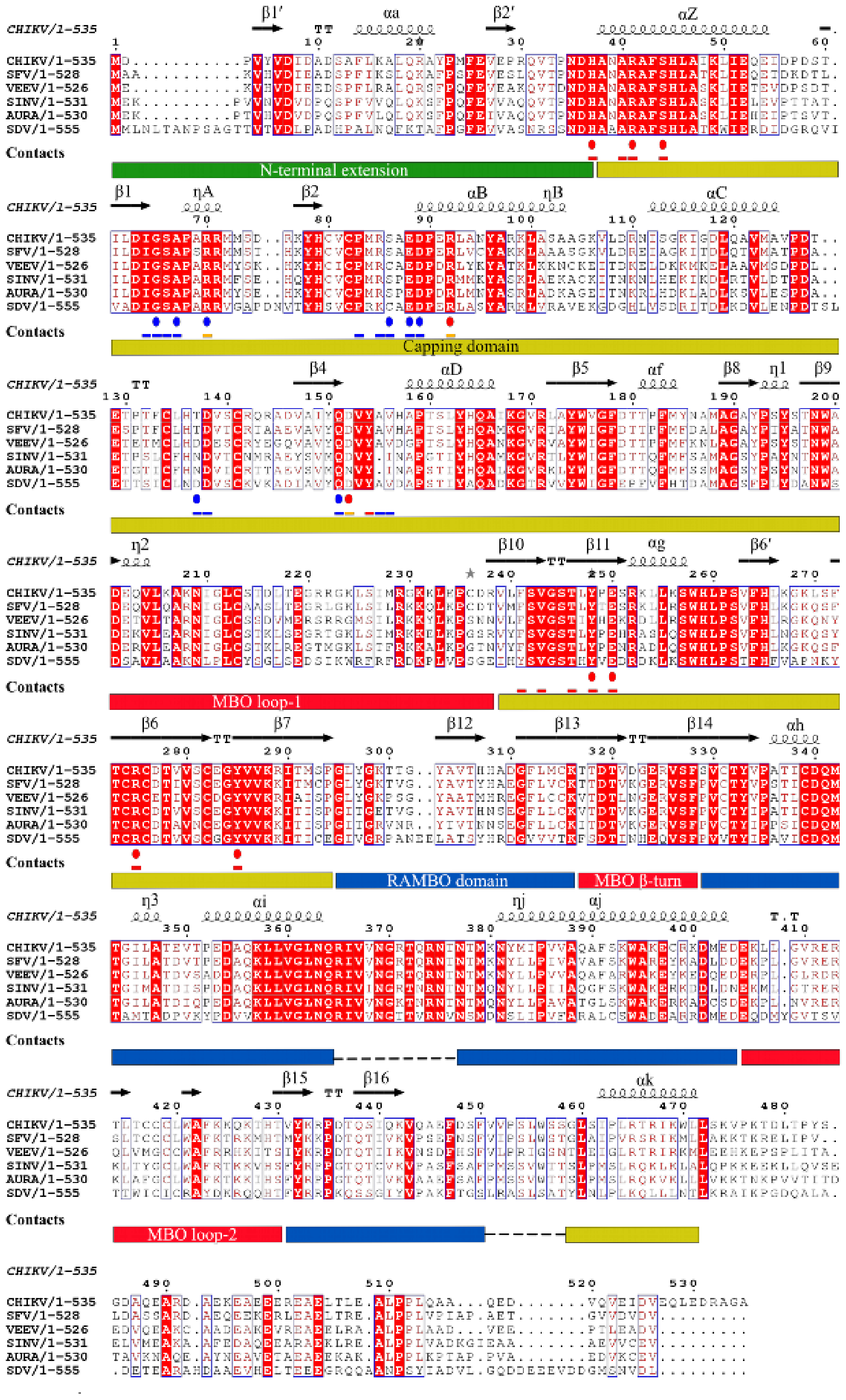
CHIKV/1-535
SFV/1-528
VEEV/1-526
SINV/1-531
AURA/1-530
SDV/1-555
Contacts
N-terminal extension
Capping domain
MBO loop-1
RAMBO domain
MBO β-turn
MBO loop-2

**Figure S2. Sequence alignment of alphavirus nsP1 proteins.** The sequence alignment of nsP1 sequences of CHIKV (UniProt: Q5XXP4), Semliki Forest virus (SFV) (UniProt: P08411), Venezuelan equine encephalitis virus (VEEV) (UniProt: P27282), sindbis virus (SINV) (UniProt: P03317), aura virus (UniProt: Q86924) and salmonid sleeping disease virus (SDV) (UniProt: Q8QL53) is shown. The secondary structural elements present in the nsP1 structure are indicated at the top of the alignment. Below the sequences, the residues that form VdW contacts to GTP, $m^7$GTP or the covalently linked $m^7$GMP are highlighted by red lines, and those forming hydrogen bonds are highlighted by red spots. Residues that contact the SAM or SAH molecules are highlighted by blue lines and H-bonds by spots. At the bottom, rectangles indicate the domain boundaries of the nsP1 fold, colored and labeled as in Jones et al. 2021. Three motifs are conserved in the amino acid sequences of alphaviruses and similar viruses such hepatitis E virus, rubella virus and more(2): (i) the catalytic histidine (H37) responsible for the cap transfer; (ii) the DXG motif (in which X stands for any residue) ($^{63}$DIG$^{65}$ in CHIKV); and (iii) and the DXXR motif ($^{89}$DPER$^{92}$ in CHIKV)—
the last two define the SAM-binding site. Driven by the structural alignment of nsP1 and *E. cuniculi* MTase, we can more generally assign the DXG motif to the SAM dependent MTases motif D/EX(G)XGXGXDL (1, 3, 4) which—in alphavirus—
corresponds to $^{63}$DIGSAPXRR$^{71}$.

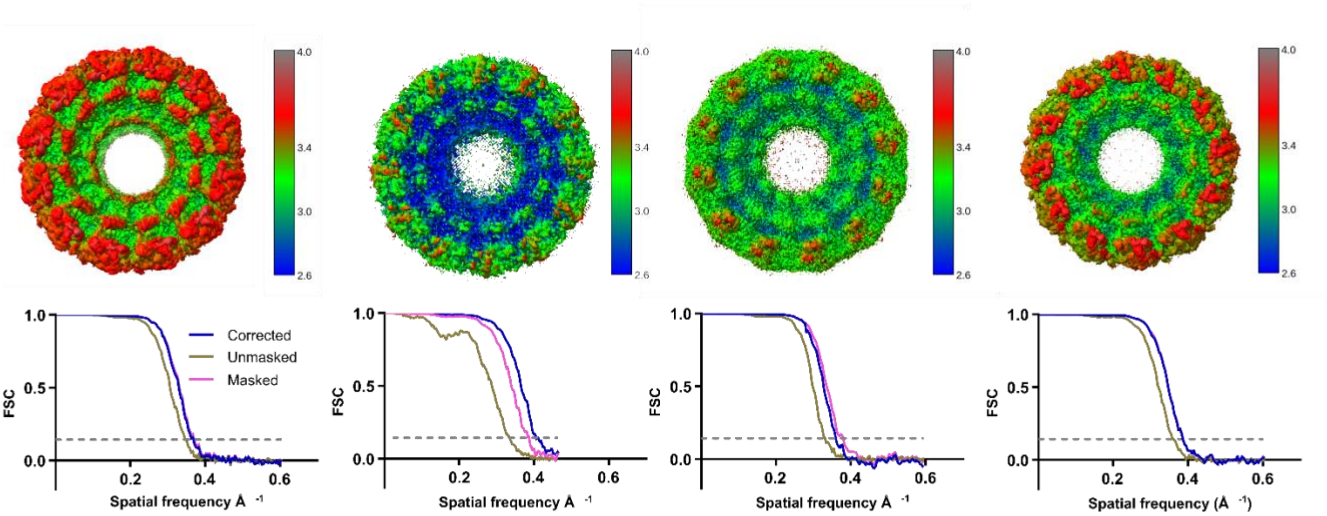


**Figure S3. Local resolution above and FSC curves below for the structures of nsP1 complexes** withnsP1+SAM (A), nsP1+GTP (B), nsP1+SAH+$m^7$GTP (C) and nsP1apo structure (D). The dashed line corresponds to the 0.143 cut-off. Resolution is significantly worse in the capping domain for the nsP1 SAM bound structure (A).

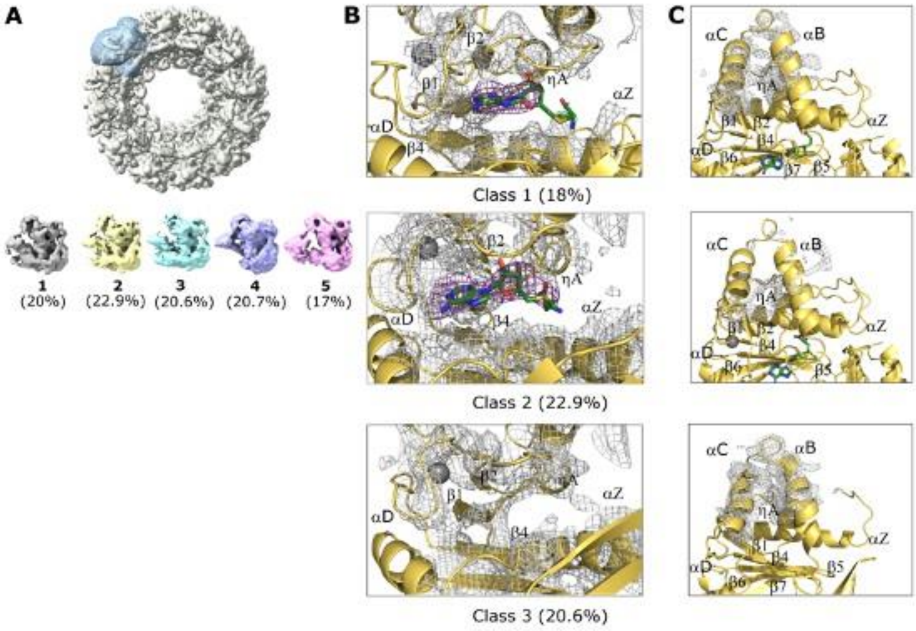


**Figure S4. Focused classification of the capping domain in the SAM bound structure.** A) Particles were symmetry expanded in Relion and signal subtracted around a soft mask focused on the capping domain from a single protomer. Focused classification was performed with varying numbers of classes. Details of the maps from classes 1, 2 and 3 are provided here, but classes 4 and 5 were too low resolution to interpret binding state properly. B) Maps showing different binding states of the SAM for class1 (purine and ribose density defined), class 2 (SAM density fully interpretable), and class 3 (site empty). Class 2 where the SAM density is fully interpretable shows a slight rotation of the SAM purine, as is observed in the SAH m$^7$GTP bound structure where the SAM is correctly positioned for the reaction. Maps are contoured at 2sigma around the binding site and density corresponding to the SAM ligand is outlined in purple. C) Corresponding density for helices $\alpha$C and $\alpha$B for the same classes. Binding of the SAM induces loss of signal in the maps for these helices, suggesting that SAM binding destabilizes this region of the structure.

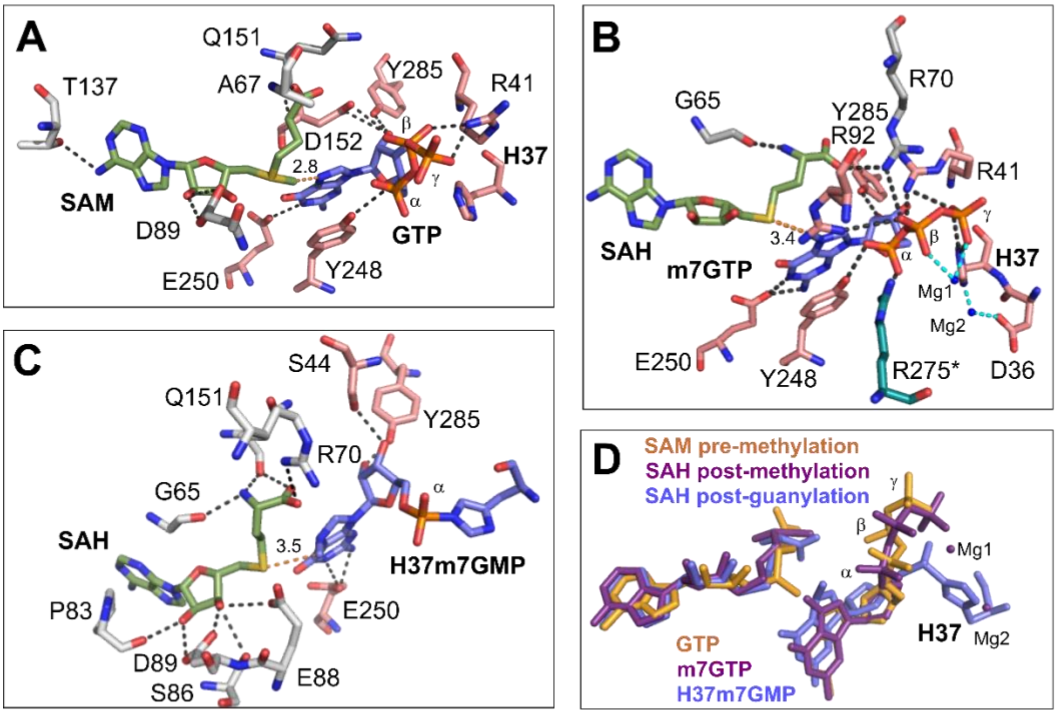


**Figure S5: Comparison of active site hydrogen bonds between the structures.** Panel A: Active site in the pre-methylation state. Residues forming contacts to the SAM ligand (green) are colored grey, whilst those contacting the GTP ligand (lilac) are in pink. The distance between the SAM methyl group and the GTP N7 (orange dashes) is indicated at 2.8Å. Panel B: Active site in the post-methylation state. Residues forming hydrogen bonds to the SAH (green) and m7GTP (lilac) ligands are colored as in A, and R257 that forms a contact from the neighbouring protomer is colored teal and indicated with an asterisk. The distance between the SAH Sulphur group and N7 methyl group of the GTP (orange dashes) increases to 3.4 Å. Magnesium ions in the active site are represented as blue spheres, and coordinating residues indicated with cyan dashes (coordinating water molecules have been removed for clarity). Panel C: Active site in the post-guanylation state. Residues are colored as in A and B, and an overall loss of hydrogen bonds formed to the m7GMP and increase in hydrogen bonds to the SAH is observed. Panel D: Superposition of the ligands between all three states (pre-methylation orange, post-methylation purple and post guanylation lilac), showing movements in the SAM/SAH ligand, guanosine ring of the GTP and a notable movement in the alpha phosphate for the H37-m7GMP bound state.

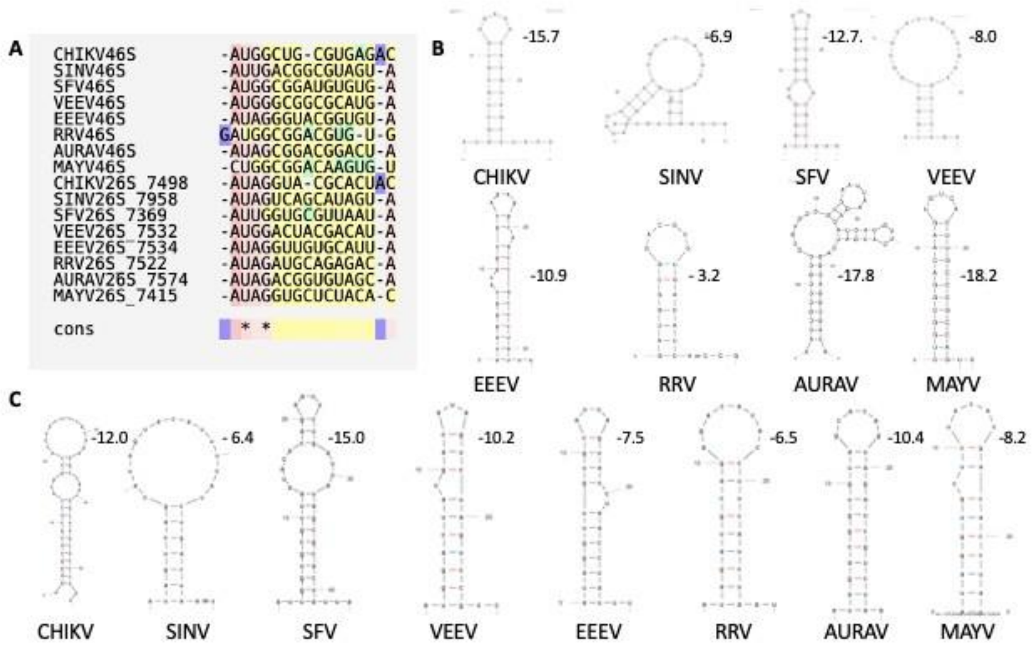


**Figure S6. Sequence and structural variation in the first structured loop (SL1) of the 5'UTR of alphaviral genomic and subgenomic RNAs.** A) Sequence alignment of the 5'UTR of genomic (46S) and subgenomic (26S) RNAs from different alphaviruses. Accession sequences used are NC_004162.2 (CHIKV), NC_001547.1 (SINV), NC_003215.1 (SFV), L01442.2 (VEEV), NC_003899.1 (EEEV), GQ433359.1 (RRV), NC_003900.1 (AURAV) and KU754168.1 (MAYV). For the 26S sequences, the initiating nucleotides for transcription of the 26S sequences are indicated after the underscore. For AURAV and MAYV, these are predicted based on alignments. Only the first 15 nucleotides are shown for clarity. Despite considerable sequence variation, the first AU and fourth G are almost universally conserved. B) Predicted secondary structures for the first stem-loop structure for 46S RNAs predicted by the mFold server, with associated predicted free energy of formation. C) Predicted secondary structures for the first stem-loop structure for 26S RNAs, as in C.

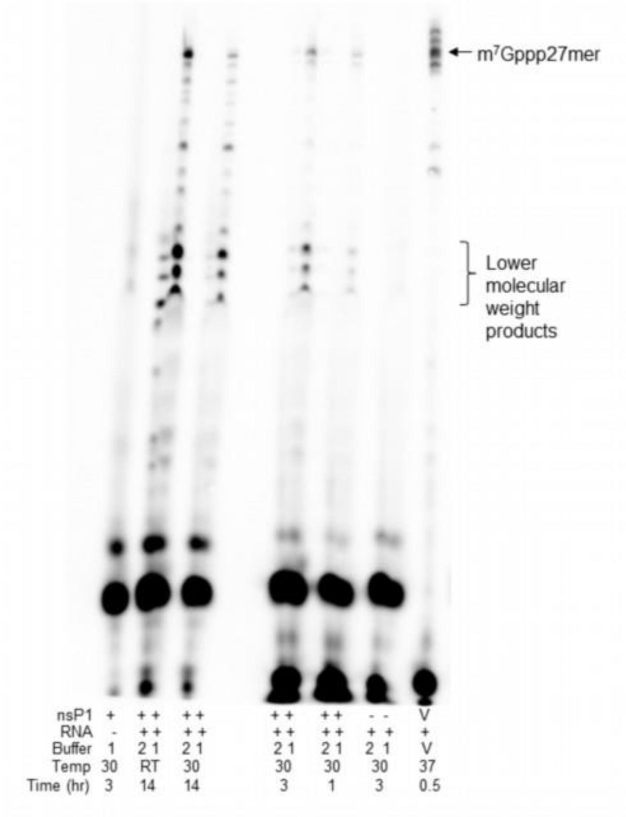


**Figure S7: Prolonged incubation of CHIKV nsP1 with the 27mer RNA leads to preferential capping of lower molecular weight products.** 2uM nsP1 was incubated with 5uM RNA 27mer in the presence of 100uM SAM and $\alpha^{32}$P labelled GTP (specific activity 0.14uCi/ul) for prolonged time periods (1-14 hours) to determine whether any capping was observed outside of the 2 hour time period typically used for experiments. Degradation products of the RNA are more strongly capped after 3 hours, and appreciable capping of the 27mer compared to the vaccinia capping enzyme control (V) is only observed with an overnight incubation. Capping activity was also compared in two buffers (buffer 1- 50mM HEPES pH 7.6, 10mM KCl, 2mM $MgCl_2$, 5mM DTT and buffer 2- 50mM tris-HCl pH 7.6, 50mM NaCl, 2mM $MgCl_2$, 5mM DTT), where cap transfer to the RNA is only observed in buffer 1.

| Sample | NsP1 + SAM | NsP1 + GTP | NsP1 + SAH + $m^7$GTP | NsP1-$m^7$GMP covalent |
|---|---|---|---|---|
| Grids | R2.2. Quantifoil Cu/Rh 300 mesh with carbon coating | R2.2. Quantifoil Cu/Rh 300 mesh with carbon coating | R2.2. Quantifoil Au 300 mesh with graphene oxide coating | R2.2. Quantifoil Au 300 mesh with graphene oxide coating |
| Vitrification method | Vitrobot | Vitrobot | Vitrobot | Vitrobot |
| Microscope | Krios CM01, ESRF | Krios CM01, ESRF | Krios CM01, ESRF | TALOS Artica, CNB Madrid |
| Magnification factor | 165,000 | 81,000 | 105,000 | 120,000 |
| Detector | K2 Summit (Gatan) | K3 Summit (Gatan) | K3 Summit (Gatan) | Falcon III |
| Recording mode | Counting | Counting | Super-resolution | Counting |
| Electron dose rate ($e^-Å^{-2}s^{-1}$) | 10.5 | 13.9 | 20.5 | 1.00 |
| Total electron dose ($e^-Å^{-2}$) | 42.4 | 42.0 | 38 | 32.0 |
| Pixel size ( Å) | 0.827 | 1.06 | 0.839 nominal, 0.42 super-resolution | 0.855 |
| Number of frames | 40 | 40 | 40 | 60 |
| Total exposure time (s) | 4 | 3.4 | 1.85 | 38 |
| Set defocus range (µm) | -1 to -2.5 | -1 to -2.5 | -1 to -2.2 | -0.8 to -2.5 |
| Number of micrographs | 4500 | 6628 | 3077 | 773 |
| Number of picked particles | 366,156 (post 2D 240,743) | 809,958 (post 2D 685,005) | 338,748 (post 2D 271,991) | 111,668 (post 2D 13,025) |
| Number of particles used for refinement | 69,057 | 95,241 | 253,384 | 10,758 |
| Symmetry imposed | D12 | C12 | C12 | C12 |
| Resolution (Å) (half-map half-map, FSC threshold 0.143) | 2.72 | 2.48 | 2.8 | 3.2 |
| RMSD bonds (Å) | 0.004 | 0.004 | 0.007 | 0.003 |
| RMDS angles (°) | 0.689 | 0.619 | 0.779 | 0.778 |
| Ramachandran favoured (%) | 97.08 | 96.82 | 97.81 | 93.32 |
| Ramachandran outliers (%) | 0.00 | 0.00 | 0.00 | 0.00 |
| Rotamer outliers (%) | 1.47 | 1.60 | 1.53 | 0.52 |
| Map-model cross correlation (CC, main chain) | 0.75 | 0.85 | 0.89 | 0.78 |
| Clashscore | 13.03 | 8.38 | 7.46 | 15.85 |
| MolProbity score | 1.92 | 1.80 | 1.59 | 2.14 |
| Map-model FSC (Å) (threshold= 0.5) | 2.8 | 2.6 | 2.7 | 3.6 |

**Table S1 Cryo-EM data collection, processing, and model refinement statistics**

| SAM/SAH binding | | | GTP/7mGTP/7mGMP-H37 binding | | |
|---|---|---|---|---|---|
| SAM | SAH-7mGTP | 7mGMP-nsP1 | GTP | SAH-7mGTP | 7mGMP-nsP1 |
| | | | | | N35 (3) |
| | | H37-7mG (15) | H37 (*:6) | H37 (3:9) | H37 covalent |
| | | | A40 (1) | A40 (4) | A40 (4) |
| | | | R41(2:11) | R41 (3:12) | R41(3) |
| | | | | | S44 (1:2) |
| I64 (2) | | I64 (1) | | | |
| G65 (11) | G65 (1:10) | G65 (1:9) | | | |
| S66 (2) | | | | | |
| A67 (1:4) | A67 (2) | | | | |
| R70 (1) | R70 (1:3) | R70 (1:3) | | R70 (3) | |
| P83 (4) | P83(7) | P83 (1:5) | | | |
| R85 (5) | R85 (2) | R85 (1) | | | |
| S86 (9) | | S86 (2:2) | | | |
| | | E88 (1:2) | | | |
| D89 (2:7) | D89 (9) | D89 (2 :8) | | | |
| R92 (5) | R92 (1) | | | R92 (1:2) | |
| T137 (3:4) | T137 (3) | T137 (3) | | | |
| D138 (8) | D138 (7) | D138 (7) | | | |
| Q151 (1:6) | Q151 (7) | Q151 (2:10) | | | |
| D152 (3) | | D152 (2) | D152 (1:13) | D152 (10) | D152 (7) |
| | | | Y154 (9) | Y154 (8) | |
| | | A155 (1) | | | |
| | V156 (2) | | | | |
| | | | F241(3) | F241 (4) | |
| | | | V243 (4) | V243 (2) | |
| | | | T246 (2) | T246 (1) | |
| | | | Y248 (1:40) | Y248 (1:26) | Y248 (13) |
| | | | E250 (2:7) | E250 (2:6) | E250 (2:3) |
| | | | | **R275 (1:3)#** | |
| | | | Y285 (2:1) | Y285(2:4) | Y285 (1:4) |
| | 7mGTP (3) | | | SAH (3) | SAH (14) |
| (7:71) | (5:57) | (10:69) | (8*:97) | (12:97) | (4:56) |

**Table S2. List of contacts of the SAM and GTP derivative ligands in the different intermediate states of the capping reaction.** For each of the structures the interacting residues are listed, indicating the number of hydrogen bonds and VdW contacts in parenthesis (H-bonds:VdW). Residues maintaining H-bonds are highlighted in red, the covalent link with H37 in highlighted in green and the residues from a neighboring nsP1 protomer in bold (R275). Contacts with the other cofactors (SAM or GTP derivatives) are also indicated in the bottom row. The total number of contacts are indicated at the bottom. *indicates possible, but not clear H-bond formation. The contacts were calculated with a distance threshold of 3.8Å for VdW contacts and 3.3Å for H-bonds using PDBSum server and LigPlot.
